# Multilayer Networks of Plasmid Genetic Similarity Reveal Potential Pathways of Gene Transmission

**DOI:** 10.1101/2022.09.08.507140

**Authors:** Julie Teresa Shapiro, Alvah Zorea, Aya Brown Kav, Vicente J. Ontiveros, Itzak Mizrahi, Shai Pilosof

## Abstract

Antimicrobial resistance (AMR) is a major threat to public health. Plasmids are principal vectors of antimicrobial resistance genes, greatly contributing to their spread and mobility across hosts. Nevertheless little is known about the dynamics of plasmid genetic exchange across animal hosts. The cow rumen ecosystem is an excellent model system because it hosts diverse plasmid communities which interact and exchange genes. Here, we use theory and methodology from network and disease ecology to investigate the potential of gene transmission between plasmids using a data-set of 21 plasmidomes from a single dairy cow population. We constructed a multilayer network based on pairwise genetic similarity between plasmids serving as a signature for past genetic exchange to identify potential routes and mechanisms of gene transmission within and between cows. The transmission network was dominated by links between cows. Modularity analysis unraveled a major cross-cow transmission pathway with additional small pathways. Plasmid functions influenced network structure: plasmids containing mobility genes were more connected; those with the same AMR genes formed their own modules. We find signatures of gene superspreading in which a few plasmids and cows are responsible for most gene exchange. An agent-based transmission model showed that a new gene invading the cow population is likely to reach all cows. Finally, we showed that link weights contain a non-random signature for the mechanisms of gene transmission allowing us to differentiate between dispersal and genetic exchange. These results provide insights into the mechanisms by which genes, including those providing AMR, spread across animal hosts.

## Introduction

Antimicrobial resistance (AMR) is a major threat to human and animal health globally [1]. Livestock may serve as a reservoir of antibiotic resistant bacteria due the widespread use of antibiotics in the agricultural sector for prophylaxis and growth promotion [2, 3]. This trend is likely to continue due to increasing demand for animal products and the intensification of livestock production globally [4]. AMR from livestock can spread into the environment, including soils and water bodies [5–8] and contaminate food products, reaching humans [9].

The spread of antibiotic resistance genes within and between species happens primarily via plasmids [10]. Plasmids are mobile genetic elements, often circular and ranging from thousands to hundreds of thousands of base-pairs long, that are able to propagate independently of their host’s chromosome. While they can be found in archaea and eukaryotes, they are most well-known in bacteria [11]. Plasmids allow bacteria to adapt to their environment by carrying accessory genes, such as those for antibiotic [12–14] or heavy metal resistance [4, 14, 15], that are beneficial under particular environmental conditions. Given the importance of plasmids to the spread of AMR [16, 17], it is important to understand how they disperse in their natural habitats between their animal hosts and interact with each other. While plasmids are one of the principal vehicles for horizontal gene transfer between bacteria, genetic exchange also occurs between plasmids themselves [11, 18]. In fact, many plasmids appear to be mosaic, incorporating genetic material from different plasmids that may be hosted by distantly related bacterial species, although this varies greatly by host taxonomy [19–21].

Livestock can be important reservoirs of plasmids containing genes for AMR [22–25]. More generally, ruminants, such as cattle, host diverse microbial communities, particularly in the rumen, which allow them to digest otherwise indigestible plant matter [26, 27]. While research on plasmids traditionally focused on those that are clinically relevant [28], advances in metagenomics and sequencing technology have allowed for the exploration of the plasmidome, all plasmids within a given sampled environment, including the bovine rumen [29, 30]. This approach has revealed that the bovine rumen plasmidome is diverse, with variation both within the rumen by space and time, and differs more between individuals compared to the bacterial microbiome [29, 30]. Based on the similarity of their open-reading frames (ORFs) to those in bacteria, the plasmids of the bovine rumen appeared most commonly associated with Firmicutes and Bacteroidetes [29, 30]. Many appear to be mosaic, including some showing evidence of cross-phyla genetic exchange [30]. Rumen plasmids are enriched for functions related to digestion and metabolism, as well as plasmid functions such as mobilization and replication but the majority of ORFs have an as-yet unknown function [29, 30]. These findings are also reported in other ecosystems such as the rat cecum where cryptic plasmids have also been found to dominate the plasmidome of [31].

Genetic exchange and dispersal may leave a signature of genetic similarity within a population, which can be detected using network analysis [32–36]. Thus far, networks of plasmid genetic similarity have primarily been used to analyze broad population structure among plasmid types and classify them into taxonomic units [33, 34, 37]. Network analyses have revealed gene sharing across geography, habitat type, and to some extent host phylogeny [34, 38, 39] identified plasmids that provide a bridge between otherwise unconnected bacterial communities [18, 40]. These uses indicate that plasmid genetic similarity networks may allow us to use signatures of past events within a plasmid population to help identify potential pathways for future transmission---an unexplored application that is critical in the context of AMR.

Because plasmids are infectious agents, disease ecology theory and methodology can prove beneficial to understanding the relationship between network structure and transmission dynamics [41–44]. Using plasmid similarity networks to understand mechanisms of gene spread is tantamount to using parasite sharing networks as a proxy for potential parasite transmission across multi-species host communities in disease ecology [42, 45], or networks that describe contacts between individuals because essentially any “transmission network” describes potential pathways for pathogen transmission and there are multiple ways to estimate these pathways [44]. In the case of gene transmission across plasmids, this approach requires that plasmid genetic information is collected within the same system. However, nearly all studies to date used plasmid sequences mined from databases and therefore originating from diverse and geographically distant environments [33, 34, 37, 39, 40]. Using plasmids from different systems is inadequate for determining pathways of genetic exchange or dispersal on finer spatial and temporal scales, for example between and within animal hosts, which are most relevant for real-life plasmid transmission. Recently, [46] used genetic similarity networks to analyze natural populations of plasmids, as opposed to sequences from databases, culturing F-type plasmids from *Enterobacterales* isolated from livestock farms (cows, pigs, or sheep) and water upstream and downstream of waste-water treatment plants. They found that the plasmid networks and communities were structured by their broad ecological niche (farm vs. water) and bacterial host genus, with a weaker effect of finer spatial scale within each environment, demonstrating the potential limits of plasmid dispersal and genetic exchange across environments and geography at ecological scales. However, in this study, samples from animals were pooled by site and thus the networks could not address transmission potential between individual hosts. Further, only cultured F-type plasmids were considered.

Here we leverage a data set of the plasmidome of the rumens of dairy cows in a single population to identify signatures of genetic exchange between plasmids and potential pathways for their dispersal between cows using a multilayer network framework. Multilayer networks are increasingly used in ecological studies [47] and allow us to explicitly consider and harness variation in network structure across multiple data layers: here, signatures of genetic similarity within and between individual cows (layers). Adopting analytical approaches and terminology from disease ecology allows us to interpret the data in light of gene transmission. We specifically aim to identify potential routes and mechanisms of gene transmission within and between cows. We find that plasmid genetic similarity networks are dominated by links between cow hosts. The transmission network is highly fragmented into clusters of plasmids (i.e., modules) with a highly heterogeneous size distribution, pointing to the dominant role of between-host transmission in shaping the genetic signature of this plasmid population. Such heterogeneity also indicates potential superspreading at the level of both plasmids and cows, whereby a few plasmids (or cows) are responsible for the majority of transmission [48]. Moreover, mobility genes were more connected, though to a limited extent. We further found that plasmids with the same AMR genes, though rare in our data set, formed independent modules. This suggests confined transmission of such genes, possibly by phylogenetic barriers or microniches created by potential spatial or chemical attributes within the ecosystem. Modeling showed that network structure determined the extent of gene transmission. By investigating the signature of genetic similarity in the network, we are able to understand how plasmids interact within and between animal hosts, providing insights into the mechanisms by which antibiotic resistance genes can spread.

## Results

### The multilayer network is dominated by interlayer, cow-to-cow connectivity

We constructed a weighted, multilayer network between plasmids sequenced by Brown Kav et al. (2020). In our multilayer network, each individual cow is a layer and nodes within each layer are plasmids (Fig 1A). Intralayer links were calculated as the genetic similarity between plasmids within a layer based on sequence alignments (Methods). A prominent feature of multilayer networks is interlayer links, which connect nodes between layers and encode ecological processes that operate between the layers. We defined the interlayer links using the same measure as intralayer links (Fig 1A), allowing us to detect signatures of gene exchange within and between cows simultaneously. Using the same definition is also advantageous because it places intra- and inter-layer processes on the same scale, avoiding a-priori biases of network metrics towards processes operating on either type of edge [36, 47, 49].

**Figure 1.**
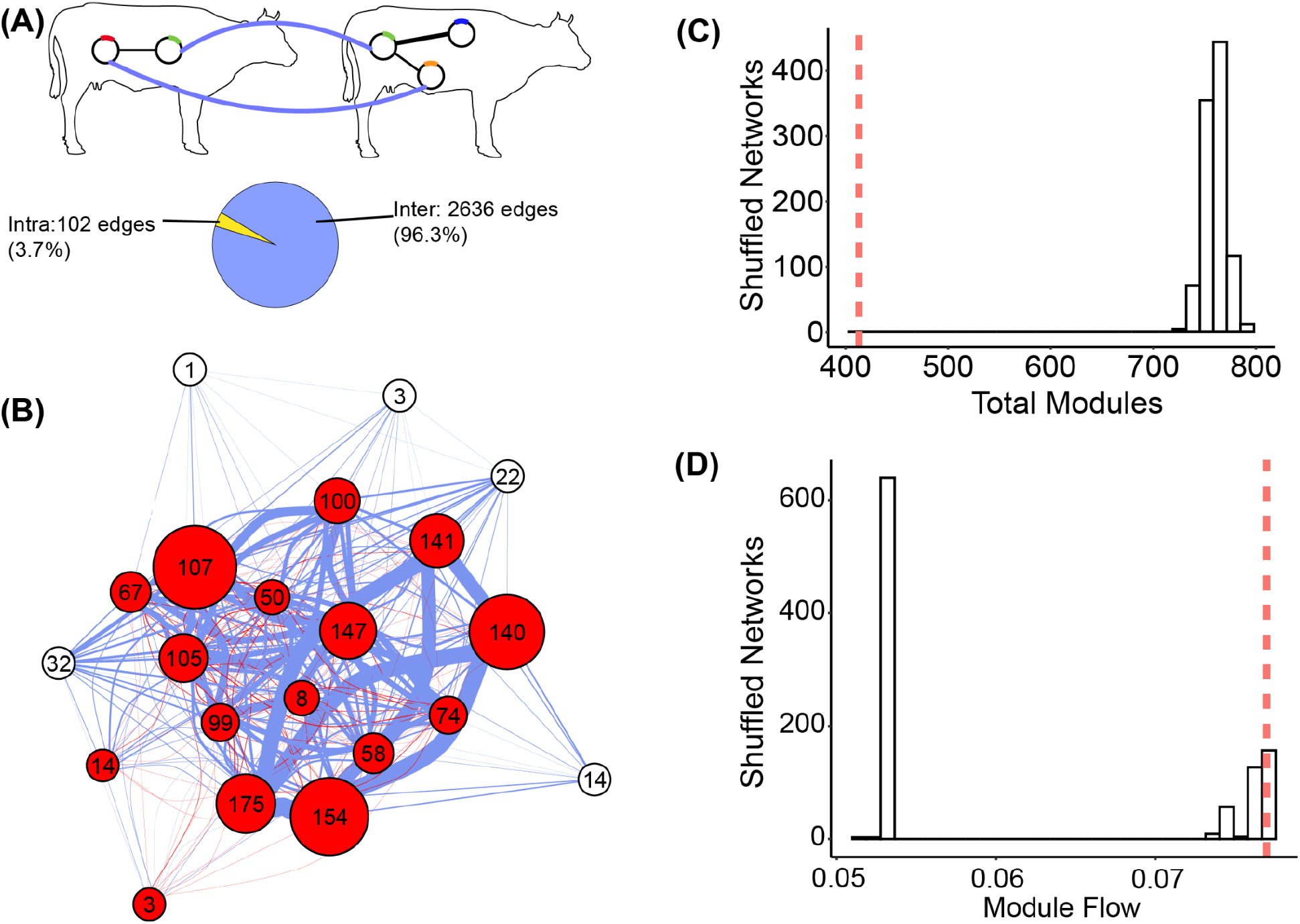
Network definition and modular structure. **(A)** Cows are layers and physical nodes are plasmids. The same plasmid can occur in different cows (e.g., the green plasmid). Intralayer (black) and interlayer (blue) links are weighted and defined as sequence similarity (see Methods). The network is dominated by interlayer edges. **(B)** A layer-perspective representation of the entire network. Circles are cows, their size is proportional to the number of intralayer edges within each cow, and their labels indicate the total number of plasmids they contain. An edge indicates that at least one interlayer edge connects two cows, and their width is proportional to the total number of interlayer edges between cows. Edges in red connect plasmids that are included in the largest module; nodes in red indicate cows that appear in the largest module. Edges in blue and nodes in white are not included in the largest module. **(C)** Comparison of the total number of modules between the observed and 1 000 shuffled networks. **(D)** Comparison of flow within the largest module of the observed and shuffled networks. In C and D the vertical dashed line indicates the value of the observed network.

Our network included 1 344 plasmids in 21 cows. Because plasmids can occur in more than one cow, the total number of nodes in the network exceeds the number of plasmids. Following standard terminology [47, 50], we define plasmids as physical nodes and a plasmid-layer combination as a state node, indicating that the same plasmid can be in a different ecological state – for instance, contributing differently to gene exchange in different cows. Our network included 1,514 state nodes and 92% of plasmids were detected in a single cow. The maximum number of cows in which a plasmid was detected was 13. The number of plasmids per cow ranged from 1 - 175, with a mean of 72. While pairs of plasmids (i.e., network links) could have multiple alignments between them, 2,728 of the 2,738 links contained a single alignment. Overall, the network was very sparse, with only 0.2% of the potential links realized.

To gauge the importance of plasmid interactions between hosts, we asked whether links were primarily within or between cows. The vast majority of edges in the network were inter-layer (Fig. 1A). However, a comparison of raw edge counts is biased by the vastly different number of possible intra- and inter-layer edges (87,002 vs. 1,058,339, respectively). Therefore, we also compared the proportion of realized intra- and interlayer links out of all possible ones (i.e., density). The density of inter-layer edges was twice as high as intra-layer edges (0.25% vs. 0.12%). Therefore, there is a much stronger signature of genetic exchange between cows compared to within them.

We tested the effect of cow identity on connectivity by comparing the observed network to 1000 randomized networks in which we shuffled the identity of cows (hereafter, ‘shuffled networks’).

We found that the observed network had a significantly higher density of intra- (p < 0.001) and inter-layer edges (p < 0.001), and a higher density (p < 0.001) than the shuffled networks (Supplementary Fig. 1), indicating that despite the sparse nature of the network, it is still more connected and with a larger contribution of inter-layer edges compared to intra-layer than expected at random. Overall, the network was dominated by interlayer connectivity, suggesting that gene exchange is more likely between plasmids from different cows than within a cow.

### Skewed plasmid contribution to gene exchange implies super-spreading

We can use the intra- and interlayer connectivity to test if certain plasmids contribute disproportionately to gene transmission and exchange by examining their degree distributions and the distribution of links to cows. Both distributions were highly skewed (skewness = 10.5 and 10.4 respectively) (Supplementary Fig. 2), with most plasmids having a few links and a few plasmids with many links to both other plasmids and cows. A skewed distribution indicates that few plasmids may be responsible for most of the transmission and interactions in the network. This pattern is in line with superspreading theory from disease ecology that consistently finds that a few hosts (here plasmids) are responsible for the majority of parasite (here an AMR gene) transmission [48, 51] (see Supplementary Information).

### The network is characterized by asymmetric and non-random pathways of gene transmission

Although the plasmid similarity network is highly sparse it may still contain major pathways of gene exchange. In epidemiology and disease ecology, areas of potential transmission in networks can be detected using algorithms of ‘community detection’, loosely referred to as modularity [41, 52] --- a mesoscale property in which parts of the network are denser compared to others [53]. We detected modules using Infomap --- an algorithm based on the movement of a random walker on the network. Infomap is designed specifically for multilayer networks and also measures the amount of flow (based on random-walk movements) contained within each node (the total flow across state nodes in the network sums to 1) [54–56]. Flow measurement is particularly suitable for our purposes because it is directly related to the idea of gene exchange [56]. Modules therefore represent high-level potential pathways of transmission.

While the network was highly fragmented with 414 modules, it was significantly less so than the shuffled networks, which had on average 83% more modules (723 - 797, mean = 760, p < 0.001) (Fig. 1B,C). As with the node-level pattern, this mesoscale topology was highly skewed: while most modules in the observed network were small, with an average of 3.3 plasmids across 3.4 cows, one exceptionally large module encompassed 12 plasmids and 16 cows (∼4 times the average number of plasmids and cows in a module). This module included 215 links, accounting for 7.9% of all the links in the network, most of which were inter-layer (n = 205) and intra-modular (n = 206). Hence, it represents a potential major pathway of transmission and genetic exchange between plasmids, particularly across different hosts (Fig. 1B). Nodes within this module encompassed 8% of the flow, which is an order of magnitude more than the mean flow per module (0.2%) (Z score = 13.3, p < 10^-5). Comparison to the largest module in each shuffled network showed that while the number of unique plasmids (physical nodes) within a module did not differ (p = 0.35), the flow in the largest module of the observed network was significantly greater than its counterpart largest modules in the shuffled networks (p = 0.003) (Fig. 1D). Therefore, more genetic exchange occurs within this large module than expected if plasmids were randomly distributed among cows. Taken together, these results point to a portion of the network where a disproportionately large amount of gene exchange occurred.

The number of modules in which cows were present was skewed (Fig. 2A) but indicated that cows share pathways of transmission. Hence, to further investigate the notion of transmission between cows, we calculated module sharing *s*_*ij*_ = |*m*_*i*_ ∩ *m*_*j*_|/*m*_*i*_. That is, the number of modules shared between pairs of cows (*m*_*i*_, *m*_*j*_), out of the total modules in cow *m*_*i*_. Module sharing was highly asymmetric, ranging from 0 - 100% (median = 14%) (Fig. 2B). As expected, cows with few modules shared all or most of them, and there was a negative correlation between the number of modules a cow hosted and the mean percent of modules shared with other cows (Kendall’s tau = -0.784698, p = 7.166e-07). Thus, a few cows may serve as hubs for a large number of interacting groups of plasmids, linking peripheral cows to transmission pathways. Nevertheless, some cows with many modules still shared a high proportion of them. For instance, four of the five cows with the greatest number of modules (125 - 164 modules) shared 53 - 69% of their modules with other cows. Thus, there are potential hotspots for plasmid transmission.

**Figure 2.**
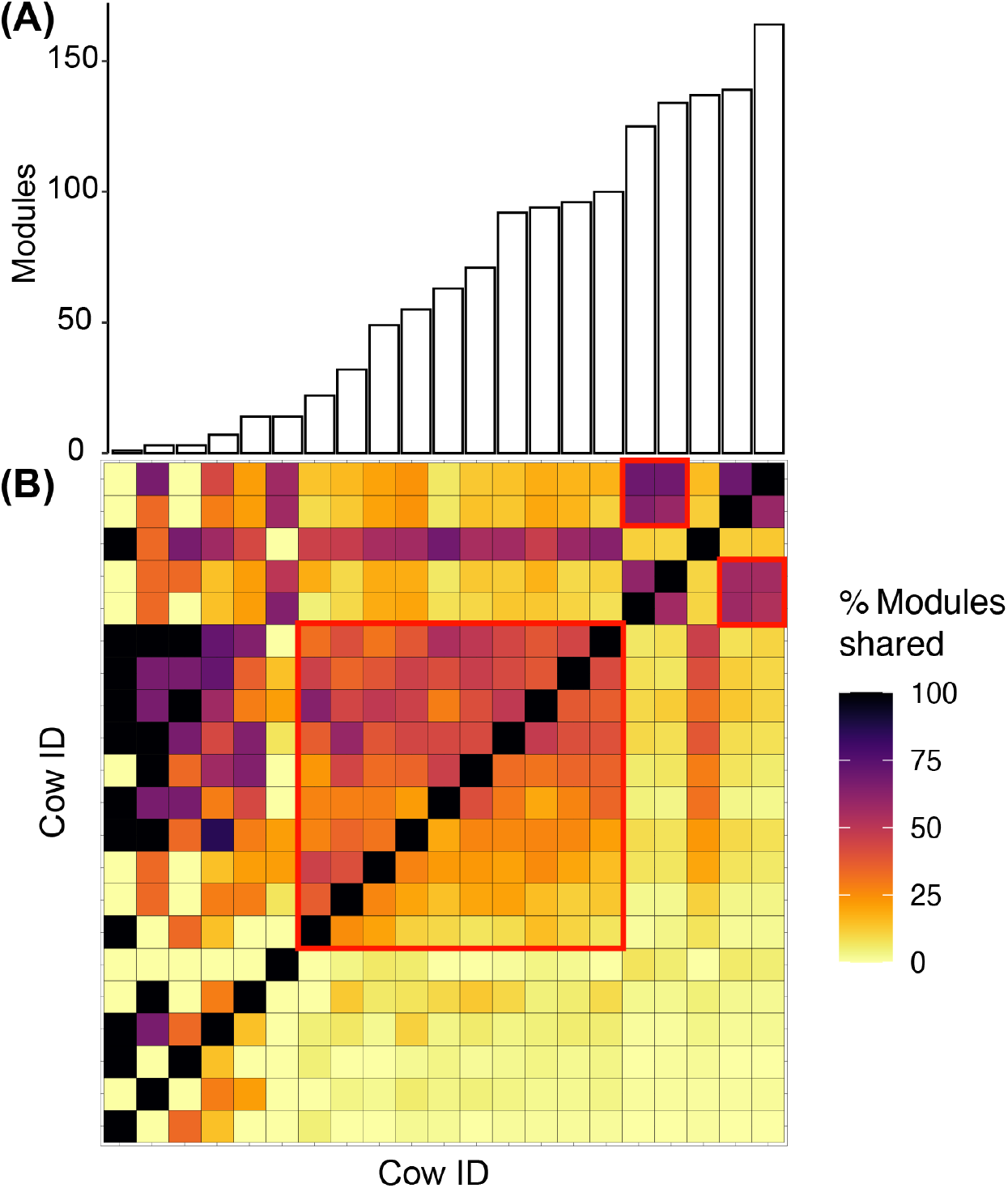
Module sharing and distribution across cows. **(A)** Barplots of the number of modules per cow. The number of modules per cow ranged from 1 - 164, with a median of 63. **(B)** The percent of modules shared between cows *i* and *j*. Each cell is calculated as: the number of modules that cow *j* shares with *i*, divided by the total number of modules (or plasmids) that *i* has. This measure results in an asymmetric matrix because the proportion is calculated with respect to one of the cows in the pair. The red square depicts an arbitrarily-chosen example for genetic exchange hotspots involving cows that share a high proportion of modules. The heatmap and barplot are ordered by the number of modules per cow, from lowest (left) to highest (right). Bars in (A) correspond to the matrix columns in (B). Cow IDs in rows and columns are the same.

The percentage of shared modules between cows was much higher than the percentage of shared plasmids, which ranged from 0 - 66.6% with a median of only 0.71% (Supplementary Fig. 3). Nearly half (47.1%) of cow pairs shared no plasmids. While this pattern is partially an artifact of the much larger number of plasmids compared to modules, it also demonstrates that while the cows largely do not host the same plasmids, they still share transmission pathways. This observation was further supported by the low correlation between the percent of shared modules and the percent of shared plasmids (Kendall’s tau = 0.41, p < 2.2e-16).

To validate the notion of cows as transmission hubs we compared observed module sharing to that obtained in the shuffled networks. For each cow pair we calculated a z-score of module sharing as 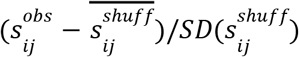. Out of the 420 possible pairs, 98 shared significantly more (z-score > 1.96) and 194 shared significantly less (z-score < -1.96) modules than the random expectation. The skew towards fewer pairs of cows sharing many modules is again consistent with potential superspreading in the network, this time at the cow level. Together, these results illustrate that the signatures of potential super-spreading are observed not only at the plasmid level, but also at the cow level.

### Simulations of plasmid-borne gene transmission between cows

We then asked how network structure might affect the spread of a hypothetical plasmid-borne gene, such as one for antimicrobial resistance, through the cow population. To address this, we constructed a stochastic agent-based model of gene transmission. In this model, the gene is initially present in a single plasmid (state node). We randomly chose 20 starting plasmids: 10 highly-connected plasmids that are a part of the largest module and 10 peripheral plasmids from the smallest modules (those containing two state nodes). In each time step, a given number of plasmids from the population encounter each other, in pairs. If one of the plasmids in the pair has the gene, the probability that the gene is transmitted to the second plasmid is equal to the genetic similarity (edge-weight) between them since research has shown that more similar plasmids are more likely to successfully horizontally transfer a gene [57]. Note that bacteria are implicit in our model and we assume that plasmid contact occurs within bacterial cells. The gene may also be lost from a plasmid with a rate depending on the level of positive selection pressure (higher selection pressure leads to lower loss rates). To determine the trade-offs between plasmid contact and selection pressure, we ran the model with low (10 plasmids in contact), intermediate (100 plasmids in contact), and high (1 000 plasmids in contact) total contact rates and accounted for selection pressure through gene loss rates: high (0 loss per capita loss rate), intermediate (0.01 per capita loss rate), and low (0.1 per capita loss rate). We ran the model for 1 000 time steps and measured the number of cows to which the AMR gene has arrived.

We found that the patterns of gene transmission were nearly identical when starting in either highly connected or peripheral plasmids (Fig. 3, Supplementary Fig. 4). In both cases, at high contact rates, the gene was able to quickly disperse to all cows in most, if not all, simulations in approximately the same amount of time at any level of selection pressure. At intermediate contact, the gene could reach all cows at high or intermediate selection pressure, although it took on average approximately 9 times as many time steps compared to the high selection pressure simulations. At low contact, it was only possible for the gene to reach all cows at high selection pressure but this only occurred in a small percent of simulations (14 - 15%) and when it did, required approximately 900 time steps on average (Fig. 3, Supplementary Fig. 4, Supplementary Table 2). Similar dynamics between highly connected or peripheral plasmids may stem from the fact that once a plasmid reaches a well-connected cow it spreads extremely quickly to others. For instance, even the five cows that are not a part of the largest module still contained other large modules (9 - 18 state nodes). Thus, even under low or intermediate selection pressure, between-cow module connectivity may allow genes to reach all hosts in this population rapidly via HGT if contact between plasmids is high enough. Another possible factor is the small cow population size (n = 21) studied here.

**Figure 3.**
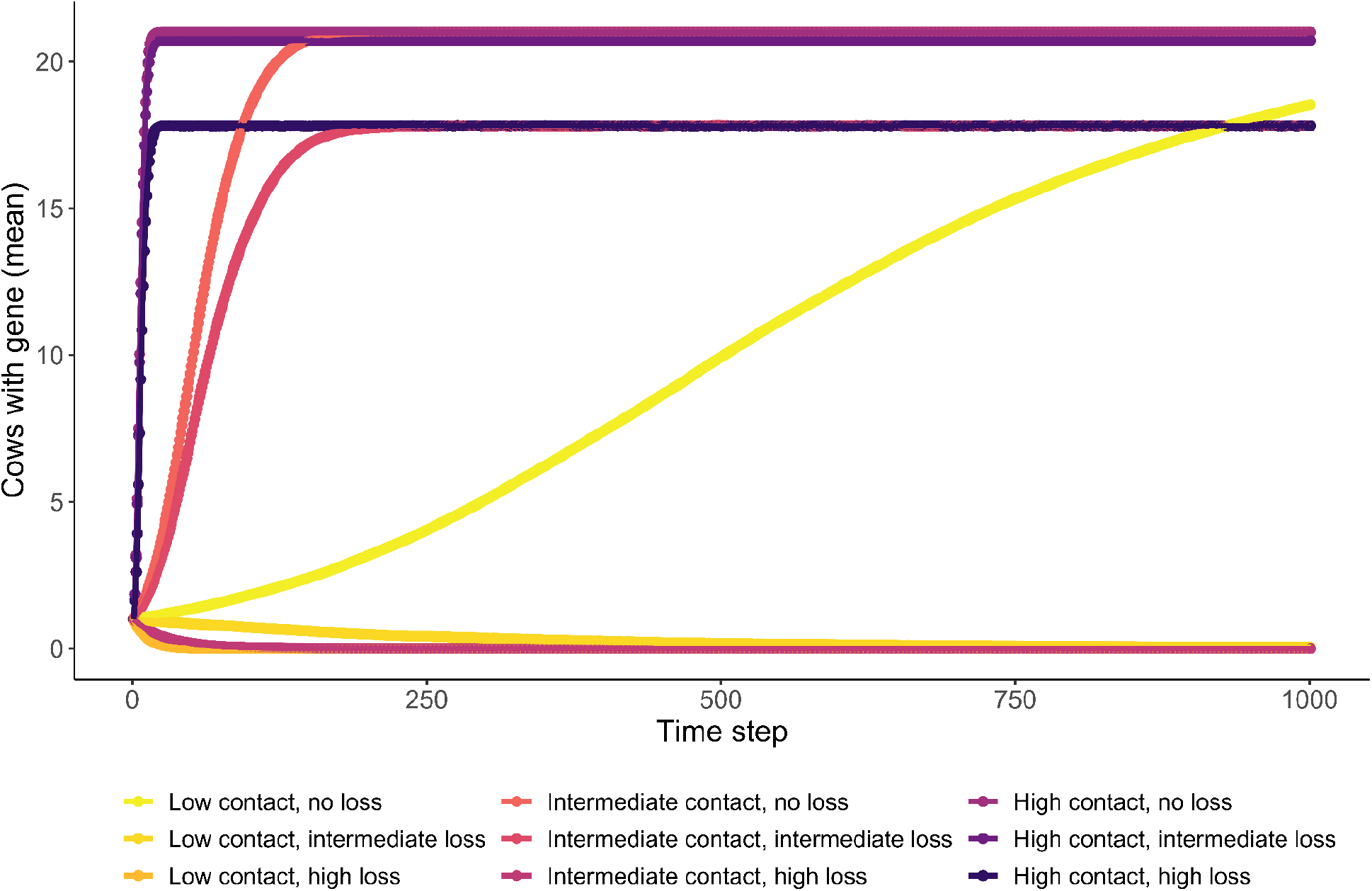
Simulated gene transmission dynamics in a cow population. Results of simulations of gene transmission among cows when the gene originates in a highly-connected plasmid. Each point is the number of cows with the gene at each time step averaged over 300 simulations per plasmid. Contact refers to the contact rate between plasmids. When plasmids encounter each other, and consequently exchange genes, at high rates, the gene is quickly transmitted to all the cow population. See Supplementary Fig. 4 for results of simulations starting in peripheral plasmids.

### Link weights are indicative of mechanisms underlying plasmid similarity

So far we have shown that there are non-random signatures of disproportionate contributions of plasmids and cows to gene exchange. However, these patterns do not suggest particular mechanisms by which gene exchange can occur. We hypothesize that such information is contained in the distribution of link weights. Specifically, plasmid dispersal should be manifested by high similarity between plasmids (very strong links) as the two nodes are essentially the same plasmid. In contrast, low link weights will be indicative of horizontal gene transfer (HGT) because they represent a small section of the plasmid’s sequence that is shared (e.g., via recombination). The distribution of edge-weights in the network displayed two abrupt breaks at 0.5 and 0.95 (Fig. 4B). We hypothesize that these breaks correspond to HGT (edge-weights < 0.5) and dispersal (edge-weights > 0.95). Between these two scenarios lies a third one---which we call “distant dispersal”---in which a plasmid disperses and then undergoes some genetic change via mutation or recombination. While it is impossible to pinpoint the particular mechanism of genetic change by pairwise genetic similarity in this scenario, such a mechanism does explain why genetic similarity lies between HGT and dispersal.

**Figure 4.**
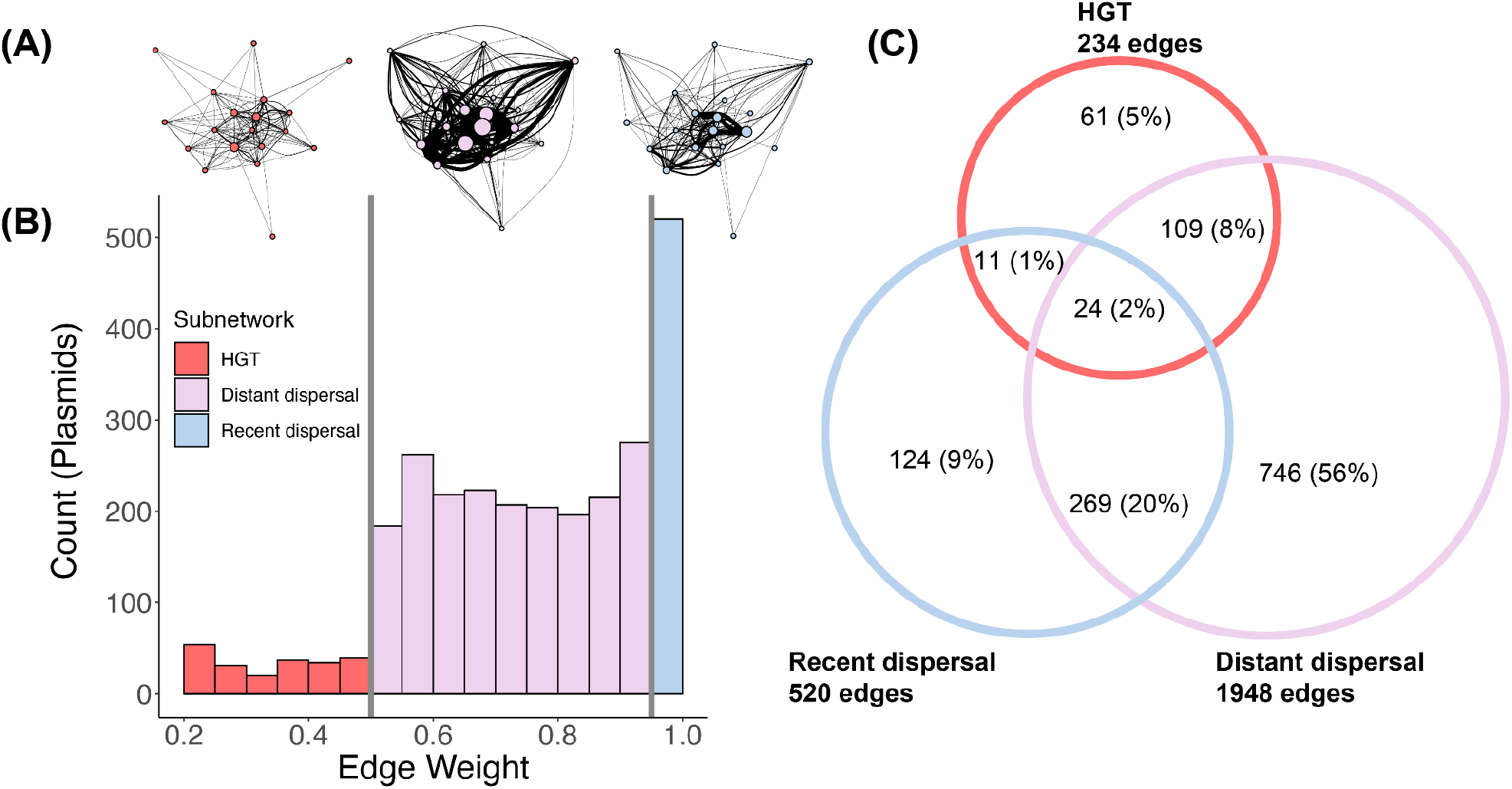
Edge-weight distribution indicating mechanisms of gene exchange. **(A)** A simplified visualization of each multilayer subnetwork (similar to Fig. 1B). Nodes are cows and their size is proportional to the number of intralayer edges within them. Edges indicate inter-layer edges between cows and their width is proportional to the total number of interlayer edges. **(B)** The distribution of edge-weights in the network have two abrupt breaks (vertical gray lines) at 0.5 and 0.95. **(C)** Venn diagram depicting the number of plasmids shared between the three subnetworks. Percentages are calculated with respect to the total number of plasmids in the data set. Circle size is illustrative of the number of plasmids and edges in the subnetwork.

The division to subnetworks represents a hypothesis that these mechanisms (HGT and dispersal) drive the observed edge-weight distribution. A first step to address this hypothesis is to test whether this distribution was non-randomly determined by sequence similarity, which is the pattern resulting from gene exchange. We resampled the alignment length between plasmid pairs without replacement 1000 times, leaving all other values the same. We then recalculated the edge-weights on the resampled data-set. The two abrupt breaks did not exist in the resampled networks. In addition, the observed edge-weight distribution was significantly different than that obtained by resampling (Kolmogorov-Smirnov test, D = 0.25, p < 0.0001), indicating that the observed distribution is likely the result of biological processes (Supplementary Fig. 5).

We then investigated each gene exchange mechanism separately by dividing the network into three multilayer subnetworks according to the link weights (Fig 4A,B). The subnetworks varied in size, with the distant dispersal subnetwork dominating in numbers of plasmids and edges (Fig. 4C). While edges could only belong to one subnetwork, plasmids could belong to multiple subnetworks (Fig. 4C). The HGT subnetwork had the highest density of intra- (0.50%) and inter-layer edges (0.78%) compared to both the recent dispersal (intra-layer = 0.38%; inter-layer = 0.22%) and distant dispersal (intra-layer = 0.11%; inter-layer = 0.25%) subnetworks. These differences in density point to the importance of HGT in driving patterns of genetic similarity seen in the network.

### Genes of mobility and antibiotic resistance affect node importance and network structure

Next, we asked whether plasmid functional traits might influence both their own importance in the network as well as the structure of the network itself. To do so, we examined the role of plasmid mobility genes, which provide the ability to transfer to new bacterial hosts. Plasmids that are conjugative (or self-transmissible) code for all the necessary proteins to transfer themselves while those that are mobilizable must use the machinery, particularly the mating pair formation (MPF) complex, of another element in the cell for transfer [58–60]. Generally, at least half of plasmids are non-mobilizable (neither conjugative nor mobilizable) [58]. To determine whether or not plasmids in our system had genes for mobilization, we used the functional annotations of the ORFs from [30]. We then measured the effect of mob genes on plasmids’ degree, module flow, and the characteristics of the modules they are found in using Mann-Whitney-Wilcoxon tests. 235 (17.4%) of plasmids in the data set had mob genes. We found that plasmids with mob genes had a significantly higher degree and greater flow, but were not found in a greater number of cows, compared to plasmids without the mob genes. Modules containing plasmids with mob genes (83 out of 414) were larger, contained more unique plasmids, encompassed more cows, and had a greater flow. Nevertheless, the magnitude of the differences in these measures was quite small (Table 1), possibly due to the low number of plasmids for which mob-related ORFs were detected.

ORFs related to antimicrobial resistance were identified in only 12 plasmids (0.9%). Eight of these were for beta-lactamase, three for tetracycline resistance, and one for penicillin-binding protein. Interestingly, plasmids with beta-lactam resistance and tetracycline resistance, respectively, each formed their own exclusive module that contained no other plasmids. The module containing the beta-lactam resistant plasmids spanned 11 different cows, making it the third largest in the network in terms of layers, perhaps indicating selective pressure promoting the persistence of these plasmids. 77% of the edges linking plasmids with beta-lactamase belonged to the distant dispersal subnetwork while the rest fell into the recent dispersal subnetwork. All the links between the three plasmids with tetracycline resistance corresponded to the distant dispersal network. Thus, within this population, plasmid dispersal appears to be the primary mechanism for the spread of plasmid-mediated antibiotic resistance between hosts.

## Discussion

A major gap in understanding the spread of AMR involves understanding how plasmid-borne AMR genes spread within and between microbial communities and animal hosts. We addressed this gap by drawing upon theory and methods from microbial and disease ecology, using multilayer networks of genetic similarity between plasmids in a population of dairy cows. While plasmid genetic similarity networks have previously been used to better classify plasmids [33, 34, 37] and identify the transmission of plasmid-borne genes across environments and geography at evolutionary scales [18, 38, 39], these scales are less relevant for transmission within animal populations. Here we used the signature left by genetic exchange between plasmids in their patterns of sequence similarity to understand plasmid transmission between hosts at a scale relevant for the transmission of plasmid-mediated antimicrobial resistance.

Further, by considering a wide range of genetic similarity between plasmids (16 - 100%), we used the similarity networks to discern the roles of HGT and dispersal, both of which play an important role in the spread of antimicrobial resistance [61, 62]. Our initial exploration of these new hypotheses showed that the distinctly segmented edge-weight distribution was non-random and thus was likely driven by biological processes. Nevertheless, the division among an HGT, recent dispersal, and distant dispersal network remains a hypothesis. Further research, via both modeling and experiments, is needed to test this hypothesis by observing rates of HGT and mutation in plasmid populations and determining under what conditions the edge-weight distribution can be reproduced.

Our network was dominated by inter-layer edges, linking plasmids in different cows. Research on plasmid-mediated antimicrobial resistance in healthcare settings have shown that plasmids often transfer between individuals via a single bacterial strain while they tend to transfer between species within the gut microbiome within individuals [61]. Thus, our results could indicate that these plasmids are transferred between cows carried by efficient microbial colonizers. However, plasmids have also been shown to disperse between human hosts independent of the transmission of bacteria [62]. Nevertheless, our plasmidome data does not allow us to include the bacterial hosts involved (see below).

Analyzing modularity in our network allowed us to identify potential pathways of transmission between hosts as hosts that share more similar plasmids may be more likely to share genes for antimicrobial resistance or even other pathogen types [42]. The fact that all plasmids with beta-lactam and tetracycline resistance were found in distinct modules supports the idea that modules represent potential pathways of transmission for genes, including those for antimicrobial resistance. Strengthening this conclusion is the fact that plasmid mobility strongly affected the modular structure. The presence of several large modules encompassing a large proportion of cows’ rumen ecosystems in this population suggests that both plasmids and the genes they host can quickly disperse through this host population. This result was supported by the dynamical model. Because the connectivity of modules between cows is high, even if a gene appears in a peripheral cow it will easily disperse to others.

The presence of mob genes affected the partitioning of the network to modules had important effects on network structure. Previous studies on plasmid networks have also found that conjugative and mobilizable plasmids are more connected than those that are non-mobilizable [39, 63]. Plasmids with mob genes may have higher degree because they can disperse more quickly between bacteria and may also encounter other plasmids more frequently, especially since mobilizable or conjugative plasmids are more likely to be found in different bacterial families while non-mobilizable plasmids are more likely to be restricted to a single species [34]. However, it is also important to consider that many small plasmids appear able to disseminate rapidly in bacterial populations and may have an as-yet unknown transfer mechanism [63].

Our results clearly show extensive dispersal of plasmids between cows, but we do not have information on the mechanism of dispersal. In healthcare settings, plasmids transfer between patients via healthcare workers or environmental reservoirs [61, 62, 64, 65]. In cows, plasmids could be transferred via saliva during grooming, especially if rumen plasmids were regurgitated during rumination or otherwise found in the mouth. Alternatively transmission may occur via the fecal-oral route as cows may rub, sniff, or lick the genital area of other individuals [66]. Transmission could also be possible via bioaerosols from feces [67]. Combining our plasmid similarity networks with social networks, using measures such as close proximity, allogrooming, or shared space use between cows [43, 66, 68] could provide further insights into how plasmids are transmitted between individuals.

We found evidence for superspreading, including the highly skewed distributions for degree at the plasmid-level and module sharing at the level of cows. Superspreading has been found in the spread of antimicrobial resistance in both humans [61, 69, 70] and cows [71]. Social networks from cows have also shown that most individuals have relatively few links to others, while a few individuals are highly connected, further indicating the potential for superspreading in the case of an outbreak [66]. Identifying patterns of superspreading are important because this can guide management and control strategies for the spread of pathogens or antimicrobial resistance and increase the effectiveness of interventions by targeting the most highly-connected individuals or groups in the network [51, 71, 72].

The transmission model we presented is a novel approach to link the structure of plasmid similarity networks with gene transmission. While plasmid similarity networks are typically used to discern processes that generated observed structures, our model can be further used to test multiple hypotheses regarding the effect of network structure on potential gene transmission, including those regarding super-spreading.

Our study has a number of limitations. First, we have not explicitly considered the plasmids’ bacterial hosts although any dispersal or interactions between plasmids would be mediated by these bacteria. Second, we do not have any measures of cow social contact. Third, our data represent only a single snapshot in time; longitudinal time-series analysis could provide further information on the dynamics of plasmid interactions and dispersal [73]. Fourth, our method of calculating plasmid similarity relies on alignments, which might not take into account rearrangements in the genomes of plasmids [33]. Finally, we only consider circular plasmids although linear plasmids also play an important role in microbial ecology [74]. Despite these limitations, our study provides the first insight into potential gene exchange via plasmid at a relevant spatio-temporal resolution.

In conclusion, genetic similarity networks provide a powerful tool for understanding transmission potential of plasmids and their genes within host populations. This approach demonstrated that within a population of dairy cows, plasmids are transmitted extensively between individuals, with potential for superspreading at the level of both plasmids and cows. Plasmid functions, particularly antimicrobial resistance and mobility, influence the network structure. The genetic similarity between plasmids in this population of cows shows signatures of both dispersal and genetic exchange, providing insights into the way plasmid-mediated antimicrobial resistance can spread across hosts.

## Methods

### Initial data processing

We used a dataset produced by [30] of plasmids sequenced from the rumens of 22 individual Israeli Holstein dairy cows housed on the same farm. The data and the methods to obtain them are described in detail in Brown Kav et al. [30]. In brief, samples of rumen fluid were obtained from each cow, DNA was extracted, amplified using phi29 polymerase, and sequenced using the Illumina paired-end protocol (Illumina GAIIX sequencer and Illumina HiSeq). Reads from each cow were first assembled into plasmid contigs using SPAdes [75]. To select only circular plasmids from the contigs, the Recycler tool [76] was used to identify closed circular sequences.

The complete plasmid dataset contained 8741 plasmid sequences. Each sequence was assigned a name based on the cow in which it was detected. We compared pairwise plasmid sequences using the BLASTn algorithm as described in Brown Kav et al. [30]. We detected identical plasmid sequences sequenced from different cows in the data-set by comparing the plasmid length, alignment length, and percent identity; identical plasmids were those with the same length and 100% identity over 100% of their length with no gaps in the alignment. We then confirmed that these plasmids were identical by aligning them in the software Geneious (v.11). A total of 314 sequences were identified as being identical to at least one other sequence in the data set and were consequently grouped in 138 of identical plasmids. Each identical plasmid was assembled from 2-5 different cows. This left 8565 unique plasmids. We did not perform any further clustering. Each unique plasmid was assigned a node id number (1-8565). Identical plasmids were assigned the same node id. Each cow was also assigned a unique layer id (1-22).

Because contigs were assembled from samples individually, we mapped the reads from each individual cow back to the full set of plasmid contigs using bbmap with the parameter “ambig” set to “all” to determine whether additional plasmids from the data set not detected in the original assembly were present in individual cows. Based on read mapping, we measured the coverage of each plasmid sequence in each cow. We considered plasmids to be present in a cow if they had 100% coverage in that cow.

### Plasmid annotations

Annotations for plasmid ORFs were previously published in [30]. These ORFs were annotated by comparing to the NCBI-NR protein database using a maximum *E*-value cut-off 10^−5^. For each ORF, the hit with the lowest E-value was chosen, unless it was a hypothetical protein, in which case we chose the next lowest E-value. If all five of the lowest E-value hits were hypothetical proteins, the ORF was annotated as hypothetical. Annotated functions were then manually curated into functional categories such as “plasmid”, “phage” and “sugar metabolism” based on their description in the database.

### Network construction

We used undirected networks because the directionality of exchange or divergence between plasmids cannot be obtained from sequence similarity alone [37, 77].

For each plasmid pair, we calculated an edge weight as:

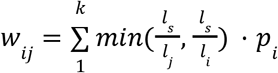

where *i* and *j* are aligned plasmids, l_s_ is the length of the alignment, l_i_ and l_j_ are the total lengths of plasmids *i* and *j, k* is the number of alignments between the plasmids and *p*_*i*_ is the percent identity between the plasmids for a given alignment. While plasmid length in the data set ranged from 2,000 - 7,297 bp (mean = 2,751 bp), aligned plasmids were generally similar in length, with 72% between 2000 - 3000 bp.) Thus, the resulting edge-weights measure the genetic similarity between plasmids.

Before constructing our networks, we performed a sensitivity analysis to determine the cut-off for plasmid length to include in our analysis. We compared the number of plasmids and alignments (total, intralayer, and interlayer) retained in the data at thresholds for plasmid length (500 to 3000 bp in increments of 500 bp) and alignment length (20% of the shorter plasmid in a given pair and 20% of the threshold). While the number of intralayer edges was relatively stable at all thresholds, we found that the number of inter-layer edges retained plateaued at a length threshold of 2000 bp while there was virtually no effect of the two alignment thresholds. Based on these results, we restricted our analyses to plasmid sequences >=2,000 bp and alignments that covered >= 20% of the length of the shortest plasmid in a pair (Supplementary Fig. 6). Minimum percent identity for alignments was >=70% following Kav Brown et al. (2020).

### Basic network metrics

We calculated the intra-, inter- and total degree of each plasmid as the number of intra-layer links, inter-layer links, and both, respectively. We tested for skewedness in the distribution of both degree centrality and layer links using the function skewedness in the package moments [78].

We calculated network density as the proportion between the total number of realized edges (defined as at least one alignment spanning ≥20% the length of the shortest plasmid in a pair) divided by the total number of potential edges. We calculated the number of potential intra- and interlayers edges, P_intra_ and P_inter_, respectively, as follows

For intra-layer edges:

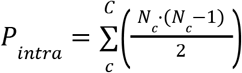

where *N*_*c*_ is the number of plasmids in cow *c*, and there are *C* cows.

For inter-layer edges:

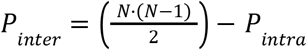

where *N* is the number of state nodes (plasmid-layer combinations) in the network: 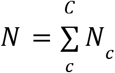.

### Shuffled networks

To determine whether the percent of realized edges and the ratio differed from random, we permuted the identity of the cows in which plasmids occur, creating 1000 shuffled networks. The algorithm conserved the distribution of edge-weights in the network as well as the number and identity of links between unique plasmids but did not constrain the number of plasmids in a cow or the number of cows a plasmid could occur in. We calculated the density and the ratio of inter- to intralayer edges of each shuffled network as described above for the observed network. We then compared the inter- and interlayer density and ratio of inter:intra-layer density in the observed network to the distribution of values for these metrics in the shuffled networks.

### Modularity using Infomap

We obtained network partitioning to modules using the infomap algorithm [54, 79] implemented in the infomapecology package [56]. In brief, infomap minimizes a function called the map equation using a modified and extended Louvain algorithm to partition the network into modules in a way that minimizes the amount of information needed to describe the movements of a random walker across the network. Modules indicate groups of nodes in the network that are more connected to each other than to other nodes. Infomap is a useful tool for analyzing this type of network because it explicitly accounts for multilayer network structure, and is computationally efficient [56]. Moreover, it is based on flow---a method particularly relevant for this study, which aims to look for signatures of gene flow. To determine whether or not the patterns of modularity and flow we observed were non-random, we applied Infomap with the same parameters as in the observed network on each of the shuffled networks (described above) and compared the distribution of modularity, module characteristics, flow, and characteristics of the largest modules in the shuffled networks to the observed network.

### Statistical analyses

When comparing measures of an observed network to that obtained from shuffled networks, we calculated p-values as the proportion of shuffled networks with values greater or lower than the observed value. This way of comparing observed to shuffled networks is a well established and common practice in network ecology [80–82].

We calculated all correlations and their p-values using the function cor.test in the stats package of program R. We used Kendall’s tau to measure all correlations due to non-normality of the data and the presence of outliers. We ran Mann-Whitney-Wilcoxon tests using the function wilcox.test in the stats package of program R with “paired” set to “false.”

### Agent-based transmission model

We used Gillespie’s direct method [83] to obtain exact stochastic simulations for our model of gene dispersal. We considered that there were two major events. First, the copy of the gene in any state node (a plasmid in a given layer) could be lost at random. Second, any pair of state nodes could be in contact and, given appropriate conditions, spread the gene from one state node to another. For simplicity, we considered that the contact rate (contacts per time unit) of all state nodes was constant. For each contact event, the algorithm selects two state nodes at random. If one of the state nodes contains the gene, it can transmit to the other state node with a probability equal to the similarity between both state nodes. Gene loss was determined by a *per capita* loss rate, and therefore for each loss event, we chose one gene for removal among the state nodes containing the gene, with equal probability. Each simulation of the model corresponds to a single realization of this stochastic process. For each of the 20 starting plasmids, we ran the model for 1 000 unitless time steps 300 times for each unique combination of contact and loss rate.

### Code

All data management and analysis were conducted in R v.4.1.1 [84].

## Acknowledgements / Funding

This work was supported by an Israel Science Foundation research grant 1281/20 to SP and a Zuckerman STEM Leadership Program postdoctoral fellowship to JTS. VJO was supported by a Margarita Salas grant funded by the Spanish Ministry of Universities and the “European Union - Next GenerationEU.” Additional funding sources included a European Research Council grant (866530) and Israel Science Foundation research grant (1947/19) to IM.

## References

1. WHO. Global Antimicrobial Resistance and Use Surveillance System (GLASS) Report: 2021. 2021. WHO.

2. Van Boeckel TP, Glennon EE, Chen D, Gilbert M, Robinson TP, Grenfell BT, et al. Reducing antimicrobial use in food animals. Science 2017; 357: 1350–1352.

3. O’Neill J. Antimicrobials in Agriculture and the Environment: Reducing Unnecessary Use and Waste. 2015. The Review on Antimicrobial Resistance.

4. Van Boeckel TP, Brower C, Gilbert M, Grenfell BT, Levin SA, Robinson TP, et al. Global trends in antimicrobial use in food animals. Proc Natl Acad Sci U S A 2015; 112: 5649–5654.

5. Managaki S, Murata A, Takada H, Tuyen BC, Chiem NH. Distribution of macrolides, sulfonamides, and trimethoprim in tropical waters: ubiquitous occurrence of veterinary antibiotics in the Mekong Delta. Environ Sci Technol 2007; 41: 8004–8010.

6. Woolhouse M, Ward M, van Bunnik B, Farrar J. Antimicrobial resistance in humans, livestock and the wider environment. Philos Trans R Soc Lond B Biol Sci 2015; 370: 20140083.

7. Noyes NR, Yang X, Linke LM, Magnuson RJ, Cook SR, Zaheer R, et al. Characterization of the resistome in manure, soil and wastewater from dairy and beef production systems. Sci Rep 2016; 6: 24645.

8. Agga GE, Cook KL, Netthisinghe AMP, Gilfillen RA, Woosley PB, Sistani KR. Persistence of antibiotic resistance genes in beef cattle backgrounding environment over two years after cessation of operation. PLoS One 2019; 14: e0212510.

9. Hudson JA, Frewer LJ, Jones G, Brereton PA, Whittingham MJ, Stewart G. The agri-food chain and antimicrobial resistance: A review. Trends Food Sci Technol 2017; 69: 131–147.

10. Gillings MR. Lateral gene transfer, bacterial genome evolution, and the Anthropocene. Ann N Y Acad Sci 2017; 1389: 20–36.

11. Rodríguez-Beltrán J, DelaFuente J, León-Sampedro R, MacLean RC, San Millán Á. Beyond horizontal gene transfer: the role of plasmids in bacterial evolution. Nat Rev Microbiol 2021; 19: 347–359.

12. Zhang T, Zhang X-X, Ye L. Plasmid metagenome reveals high levels of antibiotic resistance genes and mobile genetic elements in activated sludge. PLoS One 2011; 6: e26041.

13. Li A-D, Li L-G, Zhang T. Exploring antibiotic resistance genes and metal resistance genes in plasmid metagenomes from wastewater treatment plants. Front Microbiol 2015; 6: 1025.

14. Bukowski M, Piwowarczyk R, Madry A, Zagorski-Przybylo R, Hydzik M, Wladyka B. Prevalence of Antibiotic and Heavy Metal Resistance Determinants and Virulence-Related Genetic Elements in Plasmids of Staphylococcus aureus. Front Microbiol 2019; 10: 805.

15. Ramírez-Díaz MI, Díaz-Magaña A, Meza-Carmen V, Johnstone L, Cervantes C, Rensing C. Nucleotide sequence of Pseudomonas aeruginosa conjugative plasmid pUM505 containing virulence and heavy-metal resistance genes. Plasmid 2011; 66: 7–18.

16. Haenni M, Poirel L, Kieffer N, Châtre P, Saras E, Métayer V, et al. Co-occurrence of extended spectrum β lactamase and MCR-1 encoding genes on plasmids. Lancet Infect Dis. 2016., 16: 281–282

17. Peter S, Bosio M, Gross C, Bezdan D, Gutierrez J, Oberhettinger P, et al. Tracking of Antibiotic Resistance Transfer and Rapid Plasmid Evolution in a Hospital Setting by Nanopore Sequencing. mSphere 2020; 5.

18. Halary S, Leigh JW, Cheaib B, Lopez P, Bapteste E. Network analyses structure genetic diversity in independent genetic worlds. Proc Natl Acad Sci U S A 2010; 107: 127–132.

19. Bosi E, Fani R, Fondi M. The mosaicism of plasmids revealed by atypical genes detection and analysis. BMC Genomics 2011; 12: 403.

20. Pesesky MW, Tilley R, Beck DAC. Mosaic plasmids are abundant and unevenly distributed across prokaryotic taxa. Plasmid 2019; 102: 10–18.

21. Casjens SR, Gilcrease EB, Vujadinovic M, Mongodin EF, Luft BJ, Schutzer SE, et al. Plasmid diversity and phylogenetic consistency in the Lyme disease agent Borrelia burgdorferi. BMC Genomics 2017; 18: 165.

22. Madec J-Y, Haenni M. Antimicrobial resistance plasmid reservoir in food and food-producing animals. Plasmid 2018; 99: 72–81.

23. Ceccarelli D, Kant A, van Essen-Zandbergen A, Dierikx C, Hordijk J, Wit B, et al. Diversity of Plasmids and Genes Encoding Resistance to Extended Spectrum Cephalosporins in Commensal Escherichia coli From Dutch Livestock in 2007-2017. Front Microbiol 2019; 10: 76.

24. Auffret MD, Dewhurst RJ, Duthie C-A, Rooke JA, John Wallace R, Freeman TC, et al. The rumen microbiome as a reservoir of antimicrobial resistance and pathogenicity genes is directly affected by diet in beef cattle. Microbiome 2017; 5: 159.

25. Sabino YNV, Santana MF, Oyama LB, Santos FG, Moreira AJS, Huws SA, et al. Characterization of antibiotic resistance genes in the species of the rumen microbiota. Nat Commun 2019; 10: 5252.

26. Brown Kav A, Benhar I, Mizrahi I. Rumen Plasmids. In: Gophna U (ed). Lateral Gene Transfer in Evolution. 2013. Springer New York, New York, NY, pp 105–120.

27. Mizrahi I, Wallace RJ, Moraïs S. The rumen microbiome: balancing food security and environmental impacts. Nat Rev Microbiol 2021; 19: 553–566.

28. Dionisio F, Zilhão R, Gama JA. Interactions between plasmids and other mobile genetic elements affect their transmission and persistence. Plasmid 2019; 102: 29–36.

29. Brown Kav A, Sasson G, Jami E, Doron-Faigenboim A, Benhar I, Mizrahi I. Insights into the bovine rumen plasmidome. Proc Natl Acad Sci U S A 2012; 109: 5452–5457.

30. Brown Kav A, Rozov R, Bogumil D, Sørensen SJ, Hansen LH, Benhar I, et al. Unravelling plasmidome distribution and interaction with its hosting microbiome. Environ Microbiol 2020; 22: 32–44.

31. Jørgensen TS, Xu Z, Hansen MA, Sørensen SJ, Hansen LH. Hundreds of circular novel plasmids and DNA elements identified in a rat cecum metamobilome. PLoS One 2014; 9: e87924.

32. He Q, Pilosof S, Tiedje KE, Ruybal-Pesántez S, Artzy-Randrup Y, Baskerville EB, et al. Networks of genetic similarity reveal non-neutral processes shape strain structure in Plasmodium falciparum. Nat Commun 2018; 9: 1817.

33. Acman M, van Dorp L, Santini JM, Balloux F. Large-scale network analysis captures biological features of bacterial plasmids. Nat Commun 2020; 11: 2452.

34. Redondo-Salvo S, Fernández-López R, Ruiz R, Vielva L, de Toro M, Rocha EPC, et al. Pathways for horizontal gene transfer in bacteria revealed by a global map of their plasmids. Nat Commun 2020; 11: 3602.

35. Savary P, Foltête J-C, Moal H, Vuidel G, Garnier S. Analysing landscape effects on dispersal networks and gene flow with genetic graphs. Mol Ecol Resour 2021; 21: 1167–1185.

36. Pilosof S, He Q, Tiedje KE, Ruybal-Pesántez S, Day KP, Pascual M. Competition for hosts modulates vast antigenic diversity to generate persistent strain structure in Plasmodium falciparum. PLoS Biol 2019; 17: e3000336.

37. Brilli M, Mengoni A, Fondi M, Bazzicalupo M, Liò P, Fani R. Analysis of plasmid genes by phylogenetic profiling and visualization of homology relationships using Blast2Network. BMC Bioinformatics 2008; 9: 551.

38. Fondi M, Fani R. The horizontal flow of the plasmid resistome: clues from inter-generic similarity networks. Environ Microbiol 2010; 12: 3228–3242.

39. Tamminen M, Virta M, Fani R, Fondi M. Large-scale analysis of plasmid relationships through gene-sharing networks. Mol Biol Evol 2012; 29: 1225–1240.

40. Yamashita A, Sekizuka T, Kuroda M. Characterization of Antimicrobial Resistance Dissemination across Plasmid Communities Classified by Network Analysis. Pathogens 2014; 3: 356–376.

41. Pastor-Satorras R, Castellano C, Van Mieghem P, Vespignani A. Epidemic processes in complex networks. Rev Mod Phys 2015; 87: 925–979.

42. Pilosof S, Morand S, Krasnov BR, Nunn CL. Potential parasite transmission in multi-host networks based on parasite sharing. PLoS One 2015; 10: e0117909.

43. VanderWaal KL, Atwill ER, Isbell LA, McCowan B. Linking social and pathogen transmission networks using microbial genetics in giraffe (Giraffa camelopardalis). J Anim Ecol 2014; 83: 406–414.

44. Kauffman K, Werner CS, Titcomb G, Pender M, Rabezara JY, Herrera JP, et al. Comparing transmission potential networks based on social network surveys, close contacts and environmental overlap in rural Madagascar. J R Soc Interface 2022; 19: 20210690.

45. Dallas TA, Han BA, Nunn CL, Park AW, Stephens PR, Drake JM. Host traits associated with species roles in parasite sharing networks. Oikos 2019; 128: 23–32.

46. Matlock W, Chau KK, AbuOun M, Stubberfield E, Barker L, Kavanagh J, et al. Genomic network analysis of environmental and livestock F-type plasmid populations. ISME J 2021; 15: 2322–2335.

47. Pilosof S, Porter MA, Pascual M, Kéfi S. The multilayer nature of ecological networks. Nat Ecol Evol 2017; 1: 0101.

48. Paull SH, Song S, McClure KM, Sackett LC, Kilpatrick AM, Johnson PTJ. From superspreaders to disease hotspots: linking transmission across hosts and space. Front Ecol Environ 2012; 10: 75–82.

49. Hutchinson MC, Bramon Mora B, Pilosof S, Barner AK, Kéfi S, Thébault E, et al. Seeing the forest for the trees: Putting multilayer networks to work for community ecology. Funct Ecol 2019; 33: 206–217.

50. Kivelä M, Arenas A, Barthelemy M, Gleeson JP, Moreno Y, Porter MA. Multilayer networks. J Complex Networks 2014; 2: 203–271.

51. Lloyd-Smith JO, Schreiber SJ, Kopp PE, Getz WM. Superspreading and the effect of individual variation on disease emergence. Nature 2005; 438: 355–359.

52. Fortuna MA, Popa-Lisseanu AG, Ibáñez C, Bascompte J. The roosting spatial network of a bird-predator bat. Ecology 2009; 90: 934–944.

53. Newman MEJ, Girvan M. Finding and evaluating community structure in networks. Phys Rev E Stat Nonlin Soft Matter Phys 2004; 69: 026113.

54. Rosvall M, Bergstrom CT. Maps of random walks on complex networks reveal community structure. Proc Natl Acad Sci U S A 2008; 105: 1118–1123.

55. De Domenico M, Lancichinetti A, Arenas A, Rosvall M. Identifying Modular Flows on Multilayer Networks Reveals Highly Overlapping Organization in Interconnected Systems. Phys Rev X 2015; 5: 011027.

56. Farage C, Edler D, Eklöf A, Rosvall M, Pilosof S. Identifying flow modules in ecological networks using Infomap. Methods Ecol Evol 2021.

57. Popa O, Hazkani-Covo E, Landan G, Martin W, Dagan T. Directed networks reveal genomic barriers and DNA repair bypasses to lateral gene transfer among prokaryotes. Genome Res 2011; 21: 599–609.

58. Smillie C, Garcillán-Barcia MP, Francia MV, Rocha EPC, de la Cruz F. Mobility of plasmids. Microbiol Mol Biol Rev 2010; 74: 434–452.

59. Garcillán-Barcia Mp, Francia MV, de la Cruz F. The diversity of conjugative relaxases and its application in plasmid classification. FEMS Microbiol Rev 2009; 33: 657–687.

60. Coluzzi C, Guédon G, Devignes M-D, Ambroset C, Loux V, Lacroix T, et al. A Glimpse into the World of Integrative and Mobilizable Elements in Streptococci Reveals an Unexpected Diversity and Novel Families of Mobilization Proteins. Front Microbiol 2017; 8: 443.

61. León-Sampedro R, DelaFuente J, Díaz-Agero C, Crellen T, Musicha P, Rodríguez-Beltrán J, et al. Pervasive transmission of a carbapenem resistance plasmid in the gut microbiota of hospitalized patients. Nat Microbiol 2021; 6: 606–616.

62. Evans DR, Griffith MP, Sundermann AJ, Shutt KA, Saul MI, Mustapha MM, et al. Systematic detection of horizontal gene transfer across genera among multidrug-resistant bacteria in a single hospital. Elife 2020; 9.

63. Xue H, Cordero OX, Camas FM, Trimble W, Meyer F, Guglielmini J, et al. Eco-Evolutionary Dynamics of Episomes among Ecologically Cohesive Bacterial Populations. MBio 2015; 6: e00552–15.

64. Abe R, Oyama F, Akeda Y, Nozaki M, Hatachi T, Okamoto Y, et al. Hospital-wide outbreaks of carbapenem-resistant Enterobacteriaceae horizontally spread through a clonal plasmid harbouring blaIMP-1 in children’s hospitals in Japan. J Antimicrob Chemother 2021; 76: 3314–3317.

65. Bingen EH, Desjardins P, Arlet G, Bourgeois F, Mariani-Kurkdjian P, Lambert-Zechovsky NY, et al. Molecular epidemiology of plasmid spread among extended broad-spectrum beta-lactamase-producing Klebsiella pneumoniae isolates in a pediatric hospital. J Clin Microbiol 1993; 31: 179–184.

66. de Freslon I, Martínez-López B, Belkhiria J, Strappini A, Monti G. Use of social network analysis to improve the understanding of social behaviour in dairy cattle and its impact on disease transmission. Appl Anim Behav Sci 2019; 213: 47–54.

67. Bai H, He L-Y, Wu D-L, Gao F-Z, Zhang M, Zou H-Y, et al. Spread of airborne antibiotic resistance from animal farms to the environment: Dispersal pattern and exposure risk. Environ Int 2022; 158: 106927.

68. Boyland NK, Mlynski DT, James R, Brent LJN, Croft DP. The social network structure of a dynamic group of dairy cows: From individual to group level patterns. Appl Anim Behav Sci 2016; 174: 1–10.

69. Rocha LEC, Singh V, Esch M, Lenaerts T, Liljeros F, Thorson A. Dynamic contact networks of patients and MRSA spread in hospitals. Sci Rep 2020; 10: 9336.

70. Lerner A, Adler A, Abu-Hanna J, Cohen Percia S, Kazma Matalon M, Carmeli Y. Spread of KPC-producing carbapenem-resistant Enterobacteriaceae: the importance of super-spreaders and rectal KPC concentration. Clin Microbiol Infect 2015; 21: 470.e1–7.

71. Stein RA, Katz DE. Escherichia coli, cattle and the propagation of disease. FEMS Microbiol Lett 2017; 364.

72. Rushmore J, Caillaud D, Hall RJ, Stumpf RM, Meyers LA, Altizer S. Network-based vaccination improves prospects for disease control in wild chimpanzees. J R Soc Interface 2014; 11: 20140349.

73. Björk JR, Dasari M, Grieneisen L, Archie EA. Primate microbiomes over time: Longitudinal answers to standing questions in microbiome research. Am J Primatol 2019; 81: e22970.

74. Dib JR, Wagenknecht M, Farías ME, Meinhardt F. Strategies and approaches in plasmidome studies-uncovering plasmid diversity disregarding of linear elements? Front Microbiol 2015; 6: 463.

75. Bankevich A, Nurk S, Antipov D, Gurevich AA, Dvorkin M, Kulikov AS, et al. SPAdes: a new genome assembly algorithm and its applications to single-cell sequencing. J Comput Biol 2012; 19: 455–477.

76. Rozov R, Brown Kav A, Bogumil D, Shterzer N, Halperin E, Mizrahi I, et al. Recycler: an algorithm for detecting plasmids from de novo assembly graphs. Bioinformatics 2017; 33: 475–482.

77. Orlek A, Stoesser N, Anjum MF, Doumith M, Ellington MJ, Peto T, et al. Plasmid Classification in an Era of Whole-Genome Sequencing: Application in Studies of Antibiotic Resistance Epidemiology. Front Microbiol 2017; 8: 182.

78. Komsta F, Novomestky L. moments: Moments, cumulants, skewness, kurtosis and related tests. Comprehensive R Archive Network (CRAN). https://CRAN.R-project.org/package=moments. xAccessed 30 Apr 2022.

79. Rosvall M, Axelsson D, Bergstrom CT. The map equation. Eur Phys J Spec Top 2009; 178: 13–23.

80. Bascompte J, Jordano P, Melián CJ, Olesen JM. The nested assembly of plant--animal mutualistic networks. Proc Natl Acad Sci U S A 2003; 100: 9383–9387.

81. Vázquez DP, Poulin R, Krasnov BR, Shenbrot GI. Species abundance and the distribution of specialization in host--parasite interaction networks. J Anim Ecol 2005; 74: 946–955.

82. Fortuna MA, Stouffer DB, Olesen JM, Jordano P, Mouillot D, Krasnov BR, et al. Nestedness versus modularity in ecological networks: two sides of the same coin? J Anim Ecol 2010; 79: 811–817.

83. Gillespie DT. Exact stochastic simulation of coupled chemical reactions. J Phys Chem 1977; 81: 2340–2361.

84. R Core Team. R: A language and environment for statistical computing. R Foundation for Statistical Computing, Vienna, Austria 2021.

