## Supplementary Information for "Multilayer Networks of Plasmid Genetic Similarity Reveal Potential Pathways of Gene Transmission"

### Supplementary Text

The degree distribution of the network led us to ask if it might display scale-free behavior [1], in which the degree of nodes decreases according to the power law, following $P(k) \approx k^{-\gamma}$ where $k$ is the degree and $2 < \gamma < 3$. This topology has been observed in other networks, including those based on genetic similarity between plasmids [2]. A scale-free network is a signature of preferential attachment in which highly connected nodes become more connected and indicates little ecological or evolutionary constraints on the plasmids that are highly connected [1], as has been shown in other ecological networks [2–4]. To investigate whether our network may be scale-free, we first plotted the degree against its probability on a log-log scale (Supplementary Figure 7) and then tested the fit of the power law to the data using the function fit_power_law in the package igraph with implementation "plfit", which calculates the minimum value at which power law behavior is observed in the data [5] and tests the null hypothesis that the data come from a power law distribution using the Kolmogorov-Smirnov statistic [5, 6]. While the network as a whole did not display scale-free behavior ($\gamma = 1.68$), the tail of the distribution (degree ≥ 8 ) appeared to ($\gamma = 2.75$).

However, a fit to a particular distribution does not mean it is the best fitting distribution and therefore we then compared the fit of this tail of the distribution to the power law distribution with fits to three alternative distributions (discrete log-normal, discrete exponential, discrete Poisson) [6] using the Vuong test implemented with the function compare_distributions in the package poweRlaw in R [7]. This function uses a log-likelihood ratio as the test statistic for the null hypothesis that the two compared distributions provide equally good fits to the data. We use a two-sided p-value that indicates the probability of obtaining a log-likelihood ratio that deviates as far from zero as the observed value if the two compared distributions are equally good. A positive test statistic and $p$-value < 0.1 indicates that the power law distribution provides a better fit to the data while a negative test statistic and $p$-value > 0.1 indicates that the power law does not provide a better fit than the alternative distribution. We found that the power law did not provide a better fit than the log-normal distribution [6] (Supplementary Table S1).

### Supplementary Tables

**Supplementary Table 1.** Comparison of the fit of the power law to three alternative distributions (discrete log-normal, discrete exponential, and discrete Poisson) to the data set. The test statistic is a log-likelihood ratio. We use a two-sided p-value that indicates the probability of obtaining a log-likelihood ratio that deviates as far from zero as the observed value if the two distributions are equally good. A positive test statistic and p-value < 0.1 indicates that the power law distribution provides a better fit to the data while a negative test statistic and p-value > 0.1 indicates that the power law does not provide a better fit than the alternative distribution.

| Comparison | Log likelihood ratio for alternative distribution | p-value (two-sided) |
|---|---|---|
| Power law - Discrete log-normal | -0.44 | 0.66 |
| Power law - Discrete exponential | 1.66 | 0.10 |
| Power law - Discrete Poisson | 2.23 | 0.03 |

**Supplementary Table 2.** Results of simulation model for central and peripheral cows at each contact and loss rate showing the percent of the 300 simulations in which the gene reached all 21 cows and, in the case that all cows were reached, the mean number of time steps required.

| Plasmid | Contact rate | Loss rate | All cows with gene (%) | Time steps |
|---|---|---|---|---|
| Highly-connected | 1000 | 0 | 100 | 14.2 |
| | 1000 | 0.01 | 98.7 | 14.4 |
| | 1000 | 0.1 | 85.5 | 16.3 |
| | 100 | 0 | 100 | 127.9 |
| | 100 | 0.01 | 85.5 | 148.8 |
| | 100 | 0.1 | 0 | - |
| | 10 | 0 | 14.7 | 899.5 |
| | 10 | 0.01 | 0 | - |
| | 10 | 0.1 | 0 | - |
| Peripheral | 1000 | 0 | 100 | 14.2 |
| | 1000 | 0.01 | 98.6 | 14.4 |
| | 1000 | 0.1 | 84.5 | 16.5 |
| | 100 | 0 | 100 | 127.6 |
| | 100 | 0.01 | 83.7 | 148.3 |
| | 100 | 0.1 | 0 | - |
| | 10 | 0 | 15.6 | 903.2 |
| | 10 | 0.01 | 0 | - |
| | 10 | 0.1 | 0 | - |

**Supplementary Table 3.** Results of statistical comparisons of the number of state nodes, physical nodes, and layers per module in each subnetwork (horizontal gene transfer, recent dispersal, distant dispersal). We first compare all subnetworks together with a Kruskall-Wallis test and then perform pair-wise comparisons between each subnetwork with a Dunn test and Bonferroni correct. We specify the comparison, network metric, statistical test used, test value, and p-value. Significant p-values (< 0.05) are highlighted in bold.

| Comparison | Network Metric (per module) | Statistical test | Statistical measure | Value | p-value |
|---|---|---|---|---|---|
| Overall | State nodes | Kruskall-Wallis | Chi-squared | 26.6 | **0.000002*** |
| | Physical nodes | | | 60.8 | **6.37e-14*** |
| | Layers | | | 29.5 | **3.86e-07*** |
| | Module flow | | | 359.51 | **< 2.2e-16** |
| Recent dispersal X HGT | State nodes | Dunn test | Z-score | 3.42 | **0.002*** |
| | Physical nodes | | | 4.19 | **0.0001*** |
| | Layers | | | 3.26 | **0.003*** |
| | Module flow | | | 5.27 | **4.14e-07*** |
| Recent dispersal X Distant dispersal | State nodes | | | 4.94 | **0.000002*** |
| | Physical nodes | | | 7.73 | **3.22e-14*** |
| | Layers | | | 5.32 | **3.11e-07*** |
| | Module flow | | | -14.26 | **1.24e-45*** |
| HGT x Distant dispersal | State nodes | | | -0.26 | 1 |
| | Physical nodes | | | 0.84 | 1 |
| | Layers | | | 0.17 | 1 |
| | Module flow | | | -15.5 | **1.02e-53*** |

### Supplementary Figures

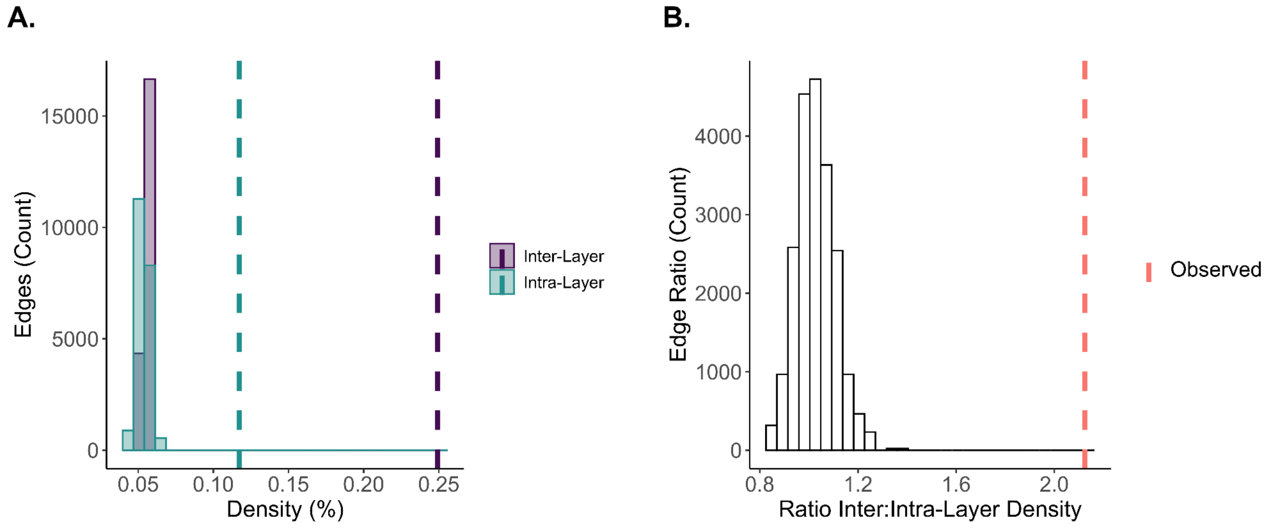


**Supplementary Figure 1.** Comparison of: (A.) density: the percent of potential inter- and intra-layer edges realized and (B.) the ratio of realized potential inter- to intra-layer edges in the observed (dashed vertical lines) and 1,000 shuffled networks (histograms). Shuffled networks were obtained by permuting layer identities.

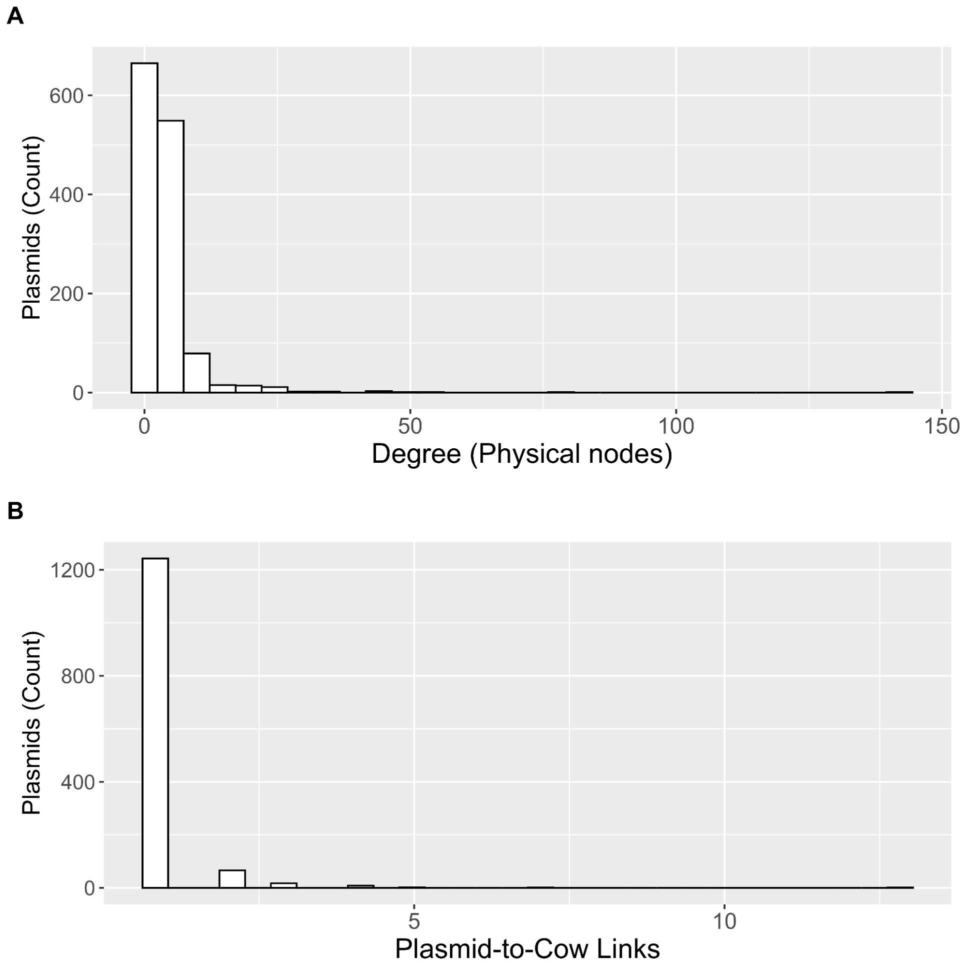


**Supplementary Figure 2.** Histogram showing the distribution of: (A) degree of plasmids (physical nodes); (B) links to cows for each plasmid.

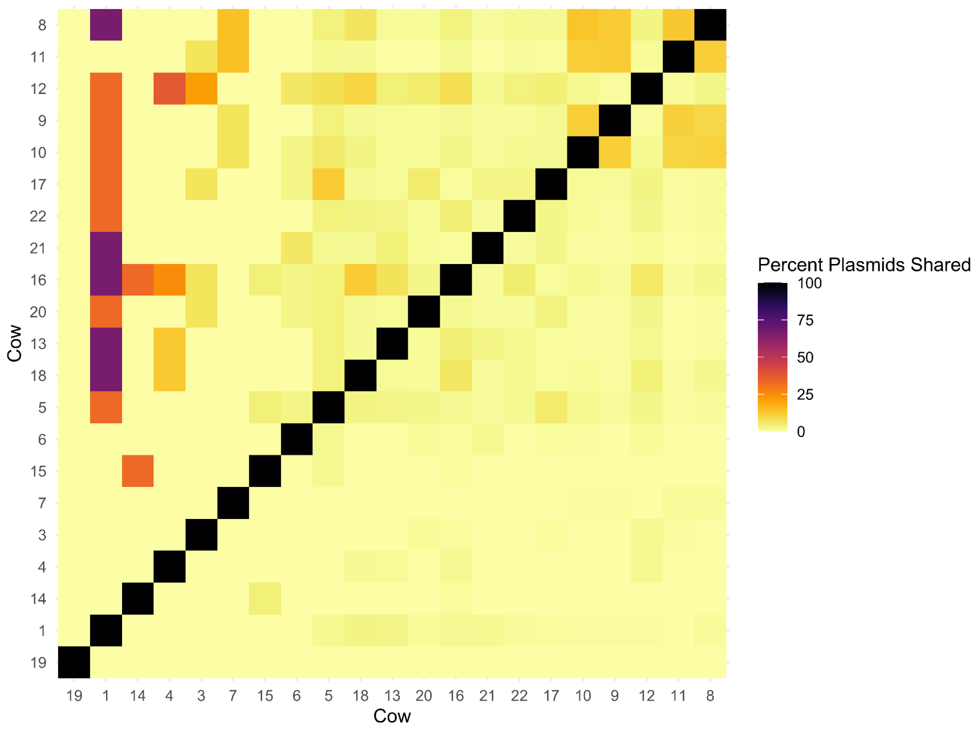


**Supplementary Figure 3.** Comparison of the percent of plasmids shared between cows *i* and *j*. The number of plasmids per cow ranged from 1 - 175, with a median of 67. The matrices are asymmetric. Each cell is calculated as: the number of plasmids that cow *j* shares with *i*, divided by the total number of plasmids that *i* has.

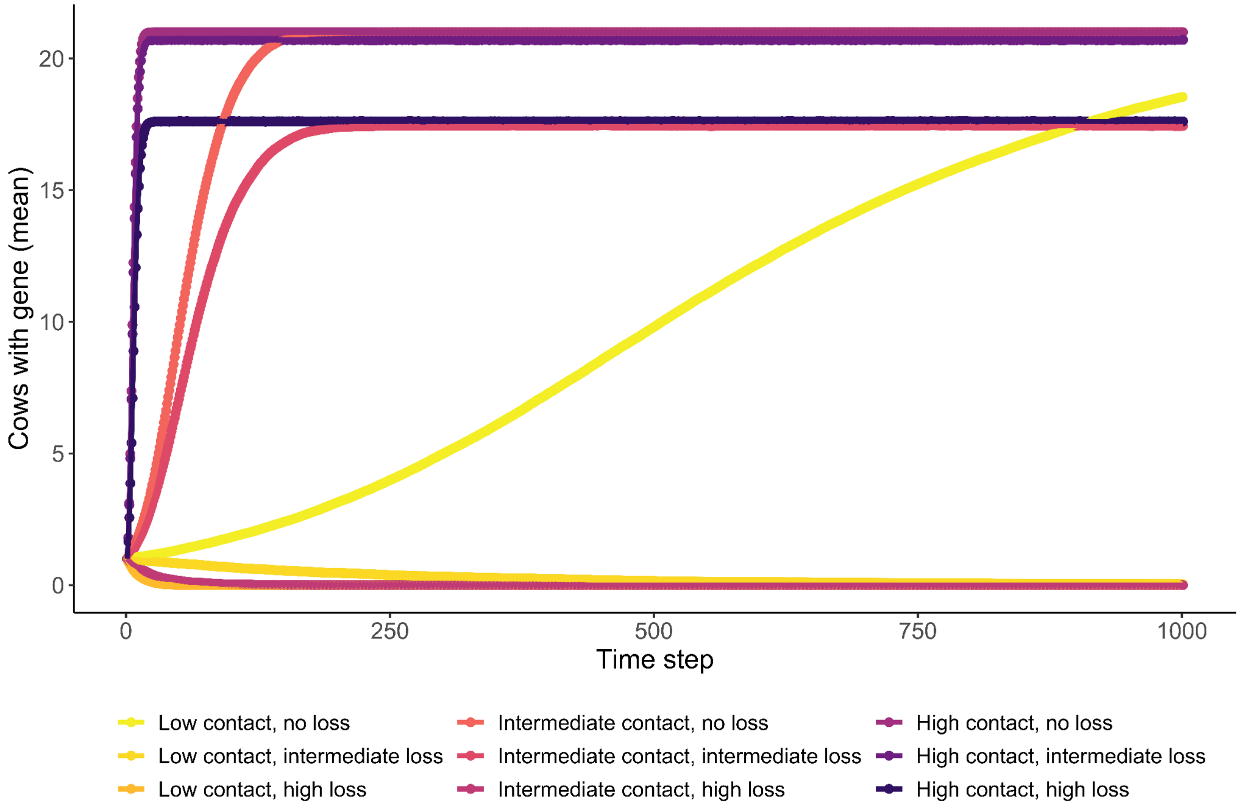


**Supplementary Figure 4. Simulated gene transmission dynamics in a cow population.** Results of simulations of gene transmission among cows when the gene originates in a peripheral plasmid. Each point is the number of cows with the gene at each time step averaged over 300 simulations per plasmid. Contact refers to the contact rate between plasmids. When plasmids encounter each other, and consequently exchange genes, at high rates, the gene is quickly transmitted to all the cow population.

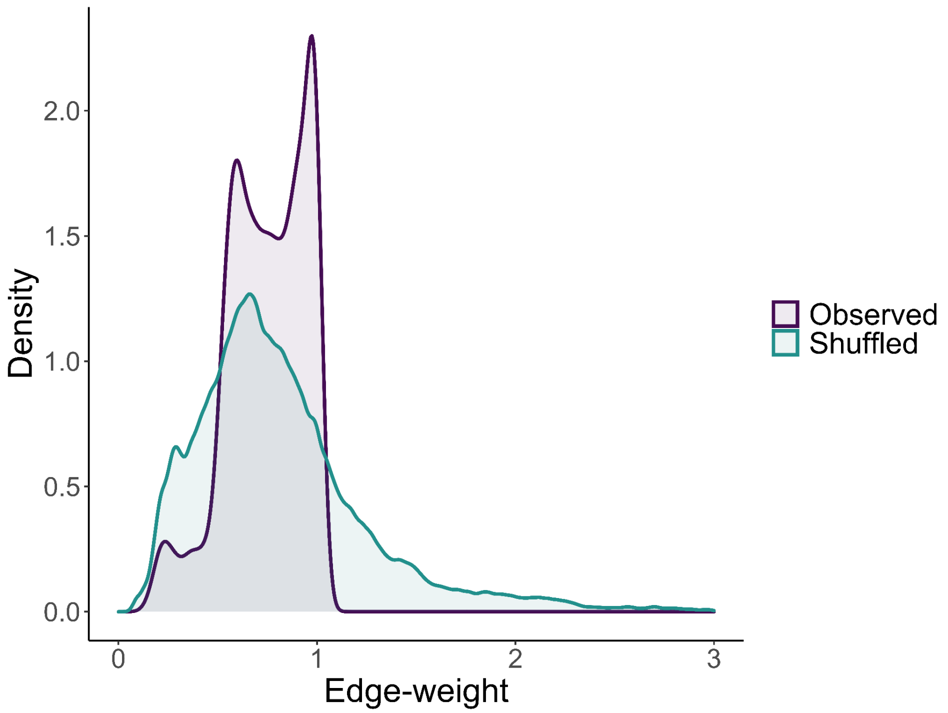


**Supplementary Figure 5.** Comparison of the distribution of edge-weights between the observed network (purple) and shuffled networks (teal). Note in shuffled networks that the alignment length could be longer than the length of either plasmid in a pair, which is not possible in the observed data.

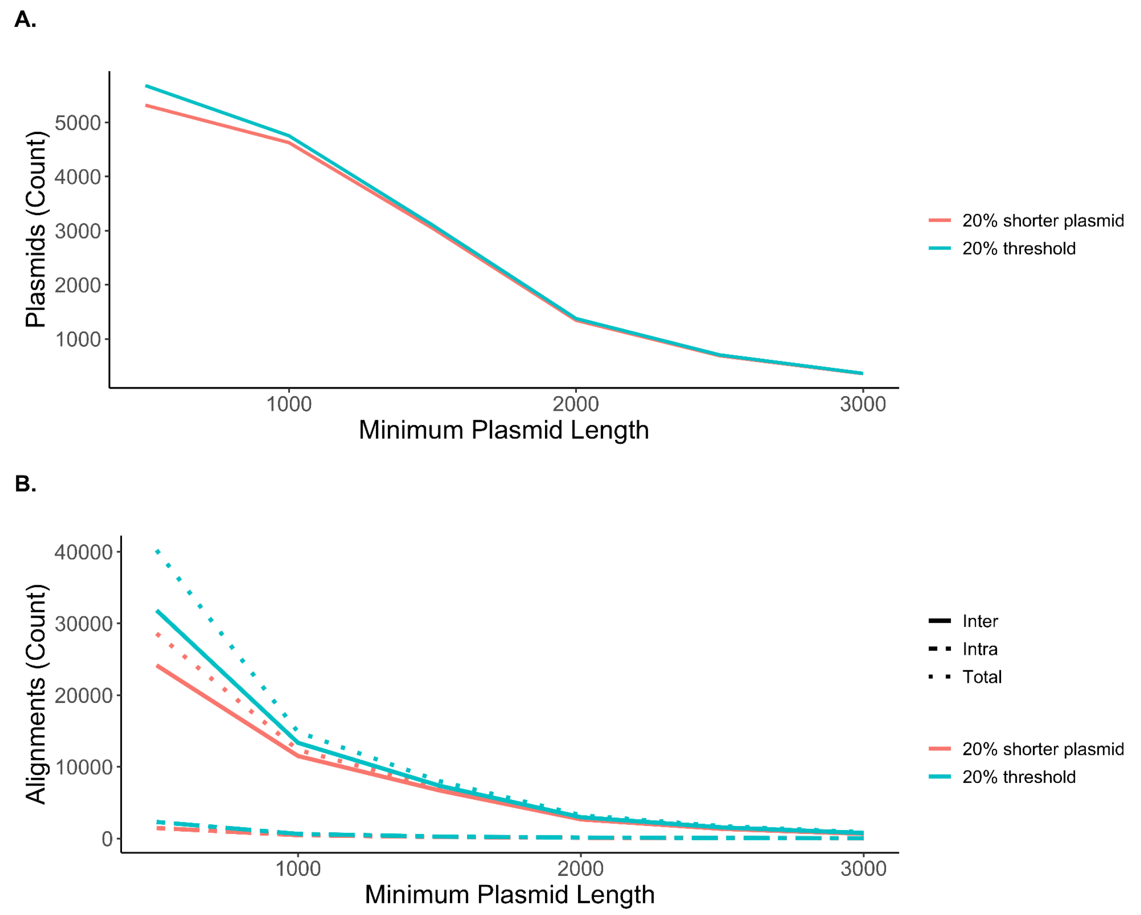


**Supplementary Figure 6.** Effect of thresholds for plasmid length and alignment length on the number of: (A.) plasmids and (B.) alignments retained in the data set.

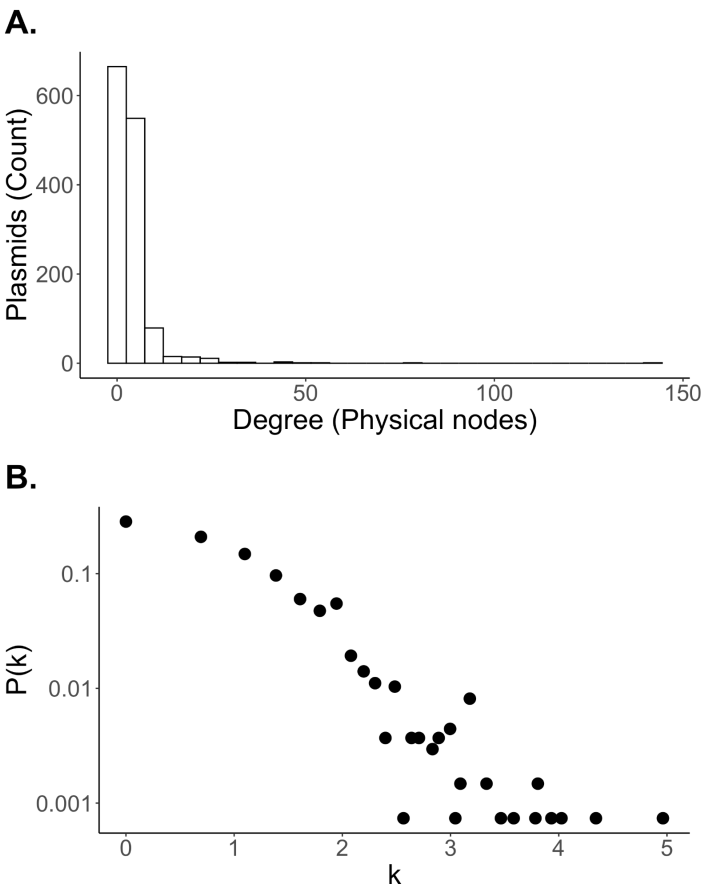


**Supplementary Figure 7.** A. Degree distribution of physical nodes. The number of links of unique plasmids (degree of physical nodes) ranged from 1 - 143 with a mean of 4.1 and median of 3. B. Log-log plot of the degree *k* and frequency of each degree P(k). The number of inter-layer links per plasmid ranged from 1 - 13 but 92.5% of plasmids had links to only one other layer.

**References**


1. Barabasi AL, Albert R. Emergence of scaling in random networks. *Science* 1999; **286**: 509–512.
2. Fondi M, Fani R. The horizontal flow of the plasmid resistome: clues from inter-generic similarity networks. *Environ Microbiol* 2010; **12**: 3228–3242.
3. Jordano P, Bascompte J, Olesen JM. Invariant properties in coevolutionary networks of plant-animal interactions. *Ecol Lett* 2002; **6**: 69–81.
4. Yamashita A, Sekizuka T, Kuroda M. Characterization of Antimicrobial Resistance Dissemination across Plasmid Communities Classified by Network Analysis. *Pathogens* 2014; **3**: 356–376.
5. Csardi G, Nepusz T. The Igraph Software Package for Complex Network Research. 2005; **Complex Systems**: 1695.
6. Clauset A, Shalizi CR, Newman MEJ. Power-Law Distributions in Empirical Data. *SIAM Rev* 2009; **51**: 661–703.
7. Gillespie CS. Fitting Heavy Tailed Distributions: The poweRlaw Package. *J Stat Softw* 2015; **64**: 1–16.
